# Mapping and Inheritance analysis of a novel dominant rice male sterility mutant, *OsDMS-1*

**DOI:** 10.1101/045625

**Authors:** Kun Yang, Yun Chen, Min Shi, Richard Converse, Xin Chen, Hengqi Min, Baimei Zhao, Yi Zhang, Jun Lv

## Abstract

We found a rice dominant genetic male sterile mutant *OsDMS-1* from the tissue culture regenerated offspring of Zhonghua 11 (*japonica* rice). Compared to wild Zhonghua 11, *OsDMS-1* mutant anthers were thinner and whiter, and could not release any pollen although the glume opened normally; most of the mutant pollen was small and malformed, and could not be stained by iodine treatment; a paraffin section assay showed the degradation of *OsDMS-1* mutant tapetum was delayed, with no accumulation of starch in the mutant pollen, ultimately leading to pollen abortion. Classical genetic analysis indicated that only one dominant gene was controlling the sterility in the *OsDMS-1* mutant. However, molecular mapping suggested three loci simultaneously control male sterility in this mutant: *OsDMS-1A,* flanked by InDel markers C1D4 and C1D5 with a genetic distance of 0.15 and 0.30 cM, respectively; *OsDMS-1B,* flanked by InDel markers C2D3 and C2D10 with a genetic distance of 0.44 and 0.88 cM, respectively; *OsDMS-1C,* flanked by InDel markers 0315 and C3D3 with a genetic distance of 0.44 and 0.88 cM, respectively. Molecular mapping disagreed with classical genetic analysis about the number of controlling genes in the *OsDMS-1* mutant, indicating a novel mechanism underlying sterility in *OsDMS-1*. We present two hypotheses to explain this novel inheritance behavior: one is described as Parent-Originated Loci Tying Inheritance (POLTI); or the hypothesis is described as Loci Recombination Lethal (LRL).

**Key message:** Three loci, which were localized on the chromosomes 1, 2 and 3 respectively, simultaneously control a dominant rice male sterility in this mutant: *OsDMS-1*.

## Introduction

Male sterility in plants includes cytoplasmic male sterility (CMS) and genetic male sterility (GMS). CMS results from the interaction between mitochondrial genes and their coupled nuclear genes (Singh and Brown 1991). GMS is caused only by nuclear genes (Vedel *et al*. 1994) and has been sub classed into Dominant genetic male sterility (DGMS) and recessive genetic male sterility (RGMS). Dominant genetic male sterility (DGMS) has proven to be a recurring mechanism. Plant DGMS was initially characterized in potatoes by Salaman (Salaman 1910). Subsequently, there have been more than 20 cases of dominant male sterility found in 10 distinct plant species (Gong and He 2006).

The mechanism of plant DGMS is more complex than that of RGMS, particularly in the restoration of sterility. Based on inheritance analysis and molecular mapping, the DGMS sterility is generally controlled by one dominant gene. Several dominant male sterile genes have been mapped (Gong and He 2006; Li and Hong 2009). Utilizing RAPD marker and BSA analysis, the *TAL* gene of Taigu genetically male sterile wheat was linked to the OPO01_900_ locus on the short arm of chromosome 4D, with a genetic distance of 14.7cM (Sui *et al*. 2001), and the rapeseed mono-dominant GMS *21-Y161* locus was found to be linked with RAPD marker S-243_1000_ (Wang *et al*. 2003), while the dominant sterility gene *Shaan-GMS* in *Brassica napus* L. was linked with BA1102_500_ (Hu *et al*. 2003). Using SRAP and SSR markers, and BSA analysis, Zhang et al. located the dominant sterility gene *CDMs399-3* in *Brassica oleracea* between the SRAP marker ENA14F-CoEm7R and the SSR marker 8C0909, with a genetic distance of 0.53cM and 2.55cM, respectively (Zhang *et al*. 2009). In rice, three dominant male sterility genes were mapped: the sterility gene *Ms-P* in Pingxiang dominant genetic male sterile rice (*PDGMSR*) was mapped to a short interval of 730 kb on chromosome 10, between two SSR markers, RM171 and RM6745 (Huang *et al*. 2007); and the *TMS* gene encoding lower temperature thermo-sensitive dominant male sterility was mapped to chromosome 6 between the SSR marker RM50 and the RFLP marker C235, with genetic distances of 12.9cM and 6.4cM, respectively (Li *et al*. 1999); utilizing SSR and InDel markers with BSA analysis, *SMS* was mapped to a 99 kb interval between InDel markers ZM30 and ZM9 on chromosome 8 (Yang *et al*. 2012). Significantly, only one gene was mapped in all of these dominant male sterility mutants which agreed with their classical inheritance analysis on the number of the loci controlling sterility.

There are two inheritance hypotheses underlying the restoration mechanism of dominant male sterility: multiple allelism, and dominant epistasis (Liu 1991; Liu 1992; Zhou and Bai 1994; He *et al*. 1999). The multiple allelism hypothesis proposes that three genes, the dominant restorer gene *Ms*^*f*^, the dominant sterility gene *Ms*, and the recessive fertility gene *ms*, can potentially occupy the same locus, with only two of these alleles appearing in each individual; their degrees of dominance are ranked as *Ms*^*f*^>*Ms*>*ms*. The dominant epistasis hypothesis proposes that there is a restorer gene, *Rf*, located in a locus distinct from the sterility gene *Ms*, with *Rf* epistatic to *Ms*. The plants containing the dominant *Ms*^+^ or *Rf* will display normal fertility regardless of the presence of the *Ms* or *ms* allele. The sterility of DGMS mutants is apparently difficult to reverse, with few DGMSs found to be restored in nature. Notable examples include a study in which extensive test crosses from 77 genetic variants of *Brassica napus* L. demonstrated that a lone Swedish cultivar, 96-803, was able to completely restore the fertility of a dominant male sterile mutant *Shaan-GMS*; an additional study in which six rice cultivars: Zigui, Penglaidao, Maiyingdao, Pingai58, IR30 and E823 were found to be restorers of the Pingxiang *Ms-P* mutant (He *et al*. 2006; Huang *et al*. 2007), with the *Rfe* restoring gene mapped to a different interval than the *Ms-P* position on chromosome 10 (Huang *et al*. 2007; Xue *et al*. 2009). These results support a dominant epistasis hypothesis.

Due to the innate difficulty in DGMS fertility restoration, homozygous mutants are rare, as a result, there exists a paucity of DGMS gene mapping and molecular characterization data. This necessitates extensive characterization of current existing mutants in order to fill this void.

Here we describe the characterization, inheritance analysis, and gene mapping of a novel DGMS mutant, *OsDMS-1,* which displays variant genetic behavior when compared to other DGMS mutants. Classical genetics suggests that a single gene controls the sterility of *OsDMS-1,* while three loci were mapped to different chromosomes in the same BF_1_ mapping population. To explain this novel disagreement between classical analysis and molecular mapping, we propose that these genes mutated in the same parent and formed a triad responsible for the sterility of *OsDMS-1*. Thus, this research provides insight into a novel DGMS mechanism.

## Materials and methods

### Plant materials

**Wild type:** Zhonghua 11 is a *japonica* variety bred by the Institute of Crop Sciences at the CAAS (Chinese Academy of Agricultural Sciences).

**Mutant type:** *OsDMS-1* was derived from the tissue-culture offspring of Zhonghua 11.

**Other fertile parents:** JH1 is an *indica* restorer bred by Southwest University; II-32B is an *indica* maintainer bred by the China National Rice Research Institute (CNRRI).

**Populations:** Three F_1_ populations including *OsDMS-1*/Zhonghua 11, *OsDMS-1*/JH1 and *OsDMS-1*/II-32B; Three BF_1_ populations including *OsDMS-1*/Zhonghua 11//Zhonghua 11, *OsDMS-1/JH1*//JH1 and *OsDMS-1*/II-32B//II-32B.

### Phenotypic observation

The fertility investigation was performed in different natural environments: Under conditions of low temperature and long day length in Beibei, Chongqing, and comparably, under conditions of high temperature and short day length in Sanya, Hainan province.

Anther protrusion and pollen release were compared by visual inspection and mature stage blooms at anthesis were photographed with an Olympus C-770 digital camera (Tokyo, Japan). Mature anthers from Zhonghua 11 and *OsDMS-1* were individually examined using a Nikon SMZ1500 stereoscope (Nikon, Tokyo, Japan) and photographed with a Nikon DS-5Mc digital camera. Anthers at bloom were crushed and stained in 1% iodine–potassium iodide (I2–KI) solution. Light microscopy was performed with a Nikon E600 microscope.

Anther development was observed on standard paraffin sections as described by Zhang (Zhang *et al*. 2008a; Zhang *et al*. 2008b) and Li (Li *et al*. 2006). Light microscopy was performed with a Nikon E600 research microscope, and photographs were taken with a Nikon DS-5Mc digital camera.

### Genetic analysis

All of the parents, F_1_, BF_1_, and the lines regenerated by fertile individuals from F_1_ and BF_1_ generations were planted during the same season. Investigation of male fertility and segregation ratios of all parents and populations was performed by visual inspection at the flowering stage. A *χ*^2^ test was used to test the goodness-of-fit.

### Molecular mapping

#### Population and sampling

BF_1_ plants from an *OsDMS-1*/JH1//JH1 population were used as a source of molecular mapping material. Equal amounts of leaf material from each of 15 random male-sterile plants were mixed, and the DNA from this mixture was extracted to construct sterile bulk. Fertile bulk was generated using a similar protocol. The fertility of all plants was recorded, and one tender leaf from each fertile and sterile plant was removed and stored for genotyping.

#### Primer design

InDel primers for the entire genome were developed based on the differences in DNA sequence between the 9311 (*indica*) and Nipponbare (*japonica*) cultivars (Shen *et al*. 2004). Remaining applied simple sequence repeat (SSR) primer sequences were downloaded from GRAMENE (http://www.gramene.org/). All the primers were synthesized by the Shanghai Sangon Biotech Corporation.

#### DNA preparation

A total of 1.5 grams of fresh flag leaves harvested at heading from either parents or populations was sampled separately, and total DNA was extracted from them utilizing a CTAB protocol(MuRRAY and Thompson 1980). DNA extraction from each BF_1_ sterile plant was performed as described (WANG *et al*. 2002) with minor modifications: (1) a portion of a tender leaf (about 1cm^2^) was cut into pieces and put into a 0.5ml centrifuge tube; (2) 100ul NaOH of 0.125M was added, boiled in a water bath for 30 seconds, and allowed to stand for a moment following removal; (3) subsequently, 50ul of 1.0M Tris-HCl (pH 8.0) was added, followed by the addition of 100ul 0.125M HCl and a 2-minute boiling water bath treatment, subsequently, the mixture was blended then allowed to stand for subsequent processing.

#### Linkage analysis

Bulk segregant analysis (BSA) (Michelmore *et al*. 1991) was adapted for linked marker selection. All InDel primers were screened against the parents and the bulks. Polymorphic primers were confirmed with 20 fertile and 20 sterile individuals. The relative chromosomal position of the *OsDMS-1* gene and its linked markers were determined by methods described by Zhang et al. (ZHANG *et al*. 2008a; ZHANG *et al*. 2008b).

## Results

### Isolation and characterization of the *OsDMS-1* mutant

In 2010, a male-sterile mutant was found among the tissue-cultured progeny of Zhonghua 11. To reproduce the mutant and analyze its inheritance, we utilized this sterile mutant as a female parent, crossing it with fertile Zhonghua 11, JH1 and II-32B. Each of three F_1_ progeny displaying fertility and sterility segregation indicated that this mutant is dominant for male sterility, thus we termed this mutant *OsDMS-1* (dominant male sterility 1). In addition to the male sterility of *OsDMS-1,* we also observed female sterility due to the fact that all of the crossing seed setting ratios were less than 15%. We also note that when compared with Zhonghua 11, *OsDMS-1* has a shorter plant height and a later heading time. Furthermore, we observed no difference in the effective tillers per plant, grain number per panicle, or 1000-grain weight between wild Zhonghua 11 and *OsDMS-1*.

To understand the impact this mutation had on flower organs and pollen development, we compared the morphology of the panicle, glume, anther, and pollen between this mutant and wild type at anthesis. No difference was observed in the flower organs between *OsDMS-1* and Zhonghua 11, and both the wild and mutant anthers protruded from the glume normally (Fig. 1A).

**Figure 1.**
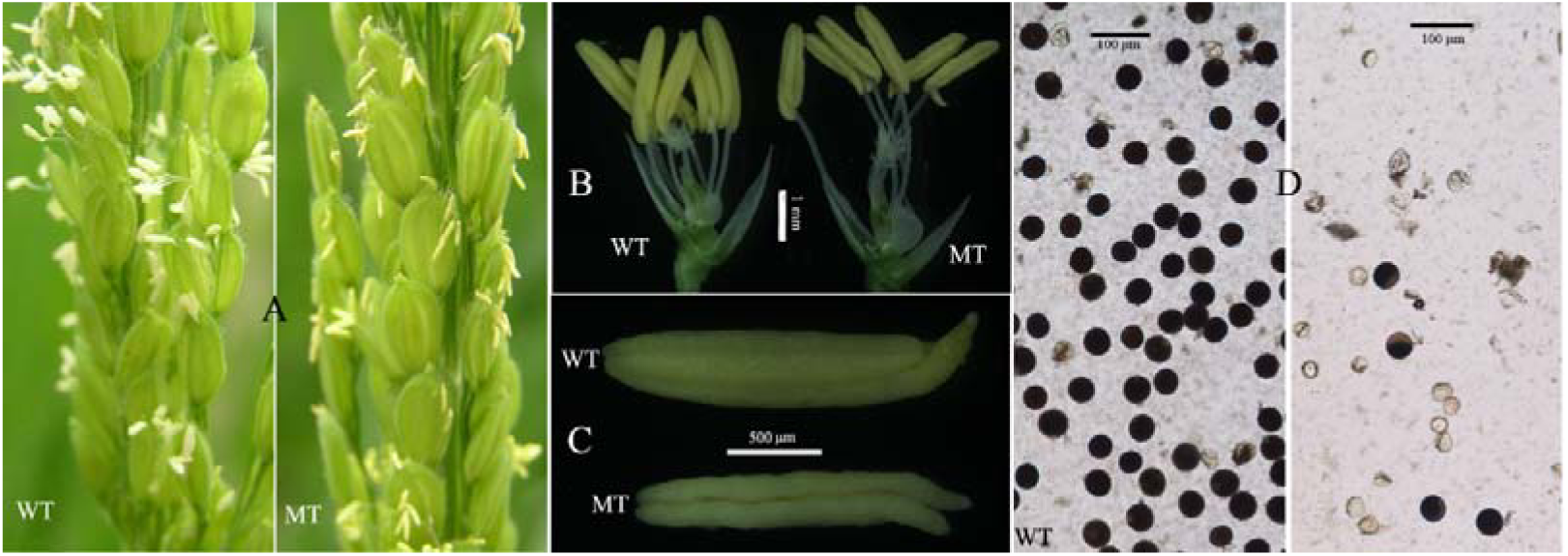
A comparison of gulme, anther and pollen appearance between Zhonghua 11 and *OsDMS-1* WT are from Zhonghua 11 (wild type). MT are from *OsDMS-1* (mutated type). D, I_2_–KI stained pollen grains showing ample black-dyed fertile pollen in the wild-type plants. only few ample pollen in the *OSDMS-1* mutant were stained by I_2_–KI.

However, obvious differences were observed post-glume opening: once protruding from the glume, the wild anthers released their pollen immediately and only left a white and empty capsule behind, while the *OsDMS-1* anthers remained un-dehiscent and inherently faint yellow, with no pollen release (Fig. 1A). Thereafter, under stereoscope, we found that the mature pre-bloom anthers in *OsDMS-1* were slender, shorter and whiter than their wild counterparts, and the surface of the mutant anther exhibited contours, unlike the smooth wild anther (Fig. 1B and C). Under the light microscope, numerous normal-sized pollen grains in crushed wild-type anthers stained with I_2_–KI solution, while fewer, smaller, malformed, unstained pollen grains were observed in *OsDMS-1* anthers (Fig. 1D).

The *OsDMS-1* mutants displayed consistent sterility in both the Chongqing low temperature/long light condition and Hainan high temperature/short light condition indicating that this phenotype was not temperature or light-sensitive.

To determine the morphological defects resident in anther tissue and pollen in the *OsDMS-1* mutant, transverse anther sections were examined. The anther development process was divided into 14 stages according as previously described (Zhang *et al*. 2011).

Using light microscopy, no detectable differences could be observed at stages 6-10 between the wild-type and mutant anthers: the pollen mother cells (PMC) were generated normally within the locule and four concentric somatic layers surrounded PMC (from the surface to interior: the epidermis, the endothecium, the middle layer and the tapetum) at stage 6 (Fig. 2A-B). The PMCs separated and moved towards the tapetal layer at stage 7 (Fig. 2C-D). Subsequently, tapetal cells became vacuolated and shrunken, with the middle layer invisible at stage 8a, while meiosis I took place and dyads were generated (Fig. 2E-F). Meiocytes underwent meiosis II continuously and generated tetrads at stage 8b (Fig. 2G-H). Free haploid microspores were released from the tetrads due to callose wall degradation, with the tapetal cells condensing at stage 9 (Fig. 2I-J). Tapetal cells gradually degraded with the round microspores increasing in volume and vacuolizing at stage 10 (Fig. 2K-L). Observable differences between *OsDMS-1* and wild Zhonghua 11 were initially noted at stage 11. The wild Tapetum degenerated almost completely into cellular debris, with only a thin epidermis of the four-layered anther wall remaining; at the same time, wild type microspores became lens-shaped binuclear pollen and began accumulating starch (Fig. 2M). However, mutant tapetal degeneration was delayed and many of the tapetal cells remained condensed and visible, with the majority of mutant microspores vacuolated like those at stage 10, without starch accumulation in the mutant pollen (Fig. 2N). At stage 12, the tapetum was invisible while the anther wall continued to thin in the wild type. At this stage, microspores became large and round, and readily stained with toluidine blue O due to the rapid accumulation of starch and other materials (Fig. 2O). Unlike wild Zhonghua 11, the *OsDMS-1* tapetum was still visible at stage 12, with no observable toluidine blue O-staining pollen due to starch accumulation failure, while the pollen became malformed (Fig. 2P). At stage 13, pollen grains gradually matured and became spherical, exhibiting deeper toluidine blue O staining due to increased starch accumulation (Fig. 2Q). The mature pollen grains were released from the anther when septum broke at stage 14 (Fig. 2S). Notably, even though the *OsDMS-1* tapetum degradation terminated, we observed minimal pollen starch accumulation at stages 13 and 14, with most of the mutant pollen small, malformed and unstained, resulting in complete sterility (Fig. 2R and T).

**Figure 2.**
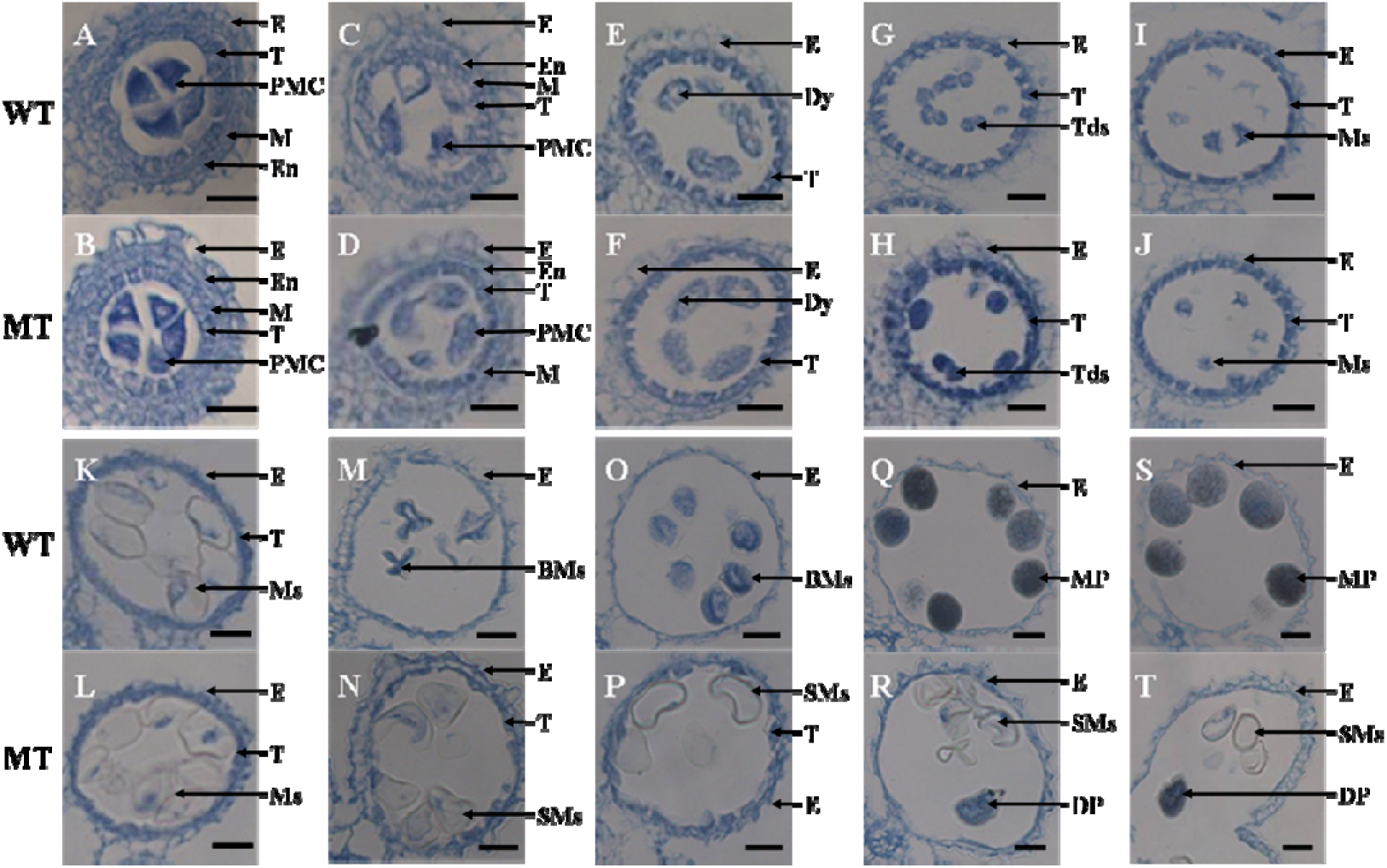
Comparison of transverse sections of wild-type (Zhonghua11) and *OsDMS-1* anthers WT, wild type. MT, mutant type. BMs, binuclear microspores. DP, dyed pollens. Dy, dyad. E, epidermis. En, endothecium. M, middle layer. MP, mature pollen. Ms, microspores. PMC, pollen mother cells. T, tapetum. Tds, tetrads. SMs, sterile microspores. Scale bar = 20 μm.

### Genetic analysis of *OsDMS-1*

To understand the inheritance of the male sterility in *OsDMS-1,* all of the parents, F_1_s, BF_1_s and the line regenerated by the fertile individuals from F_1_ and BF_1_ were grown in the same conditions in Beibei, Chongqing, from March to August, with the spikelet fertility and the segregation ratios measured at the flowering stage. A chi-square test was used to test the goodness-of-fit. The results showed: both of the F1s and BF1s demonstrated fertility segregation with a 1:1 ratio, which was confirmed by *χ*^2^ tests (Table 1). The next generation of the fertile plants from F1 and BF1 were consistently fertile among individuals from any line (data not shown). These results suggest that the male sterility of the *OsDMS-1* mutant might controlled by a single dominant sterility gene.

**Table 1.**
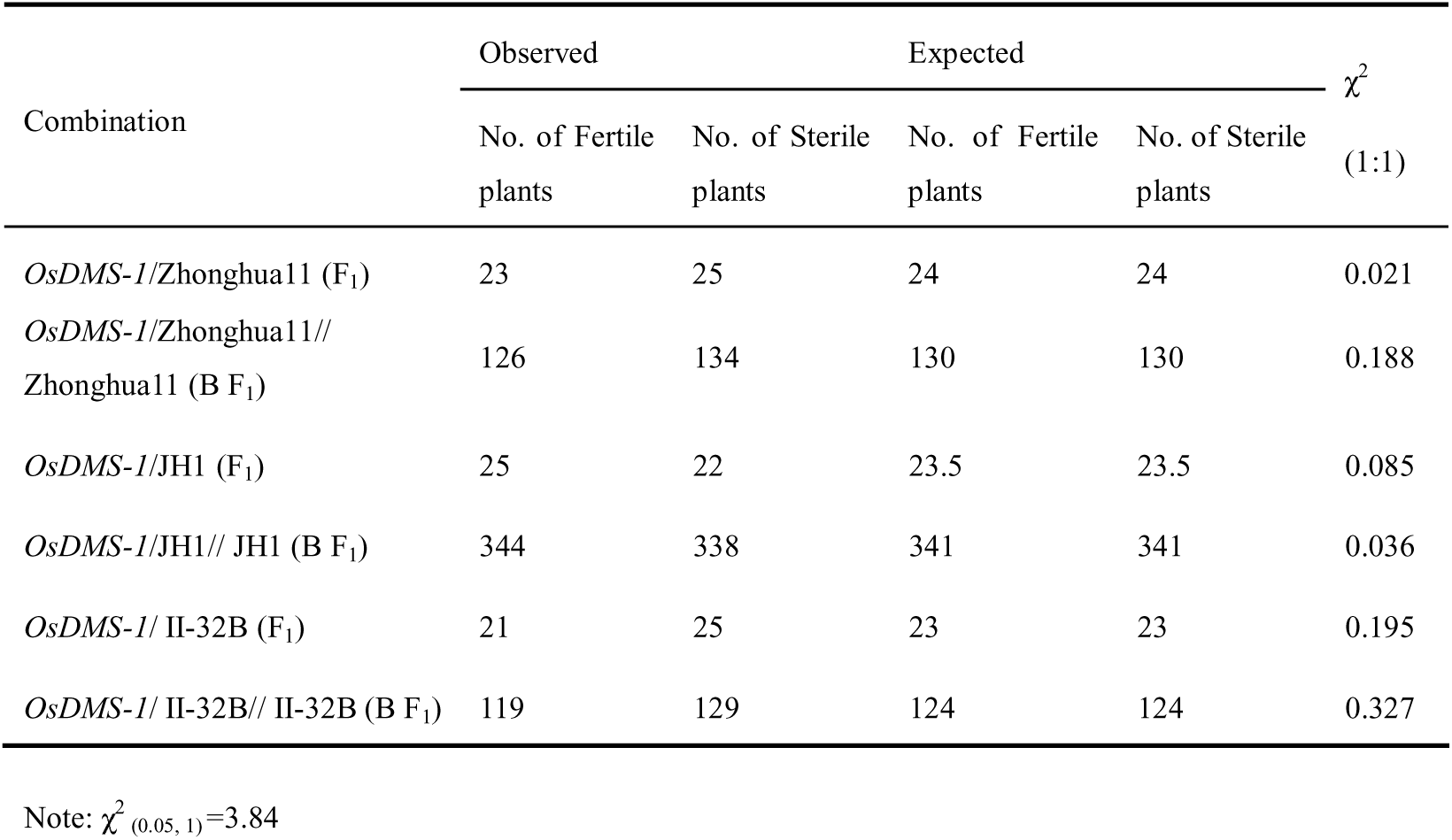
Segregation ratios of fertility in F_1_ and BF_1_ populations

| Combination | Observed | | Expected | | $\chi^2$<br>(1:1) |
| --- | --- | --- | --- | --- | --- |
|  | No. of Fertile plants | No. of Sterile plants | No. of Fertile plants | No. of Sterile plants |  |
| *OsDMS-1*/Zhonghua11 (F<sub>1</sub>) | 23 | 25 | 24 | 24 | 0.021 |
| *OsDMS-1*/Zhonghua11//<br>Zhonghua11 (B F<sub>1</sub>) | 126 | 134 | 130 | 130 | 0.188 |
| *OsDMS-1*/JH1 (F<sub>1</sub>) | 25 | 22 | 23.5 | 23.5 | 0.085 |
| *OsDMS-1*/JH1// JH1 (B F<sub>1</sub>) | 344 | 338 | 341 | 341 | 0.036 |
| *OsDMS-1*/ II-32B (F<sub>1</sub>) | 21 | 25 | 23 | 23 | 0.195 |
| *OsDMS-1*/ II-32B// II-32B (B F<sub>1</sub>) | 119 | 129 | 124 | 124 | 0.327 |

Note: $\chi^2_{(0.05, 1)}=3.84$

### Mapping the *OsDMS-1* gene

#### Linked marker screening and verification

To localize the *OsDMS-1* gene to a single chromosome, 137 InDel primer pairs evenly distributed throughout the 12 chromosomes were used to screen parents and bulks. Four primer pairs on chromosome 1, two primer pairs on chromosome 2, and four primer pairs on chromosome 3 were found to amplify polymorphic products between both parents and bulks (Table 2, bold font).

**Table 2.**
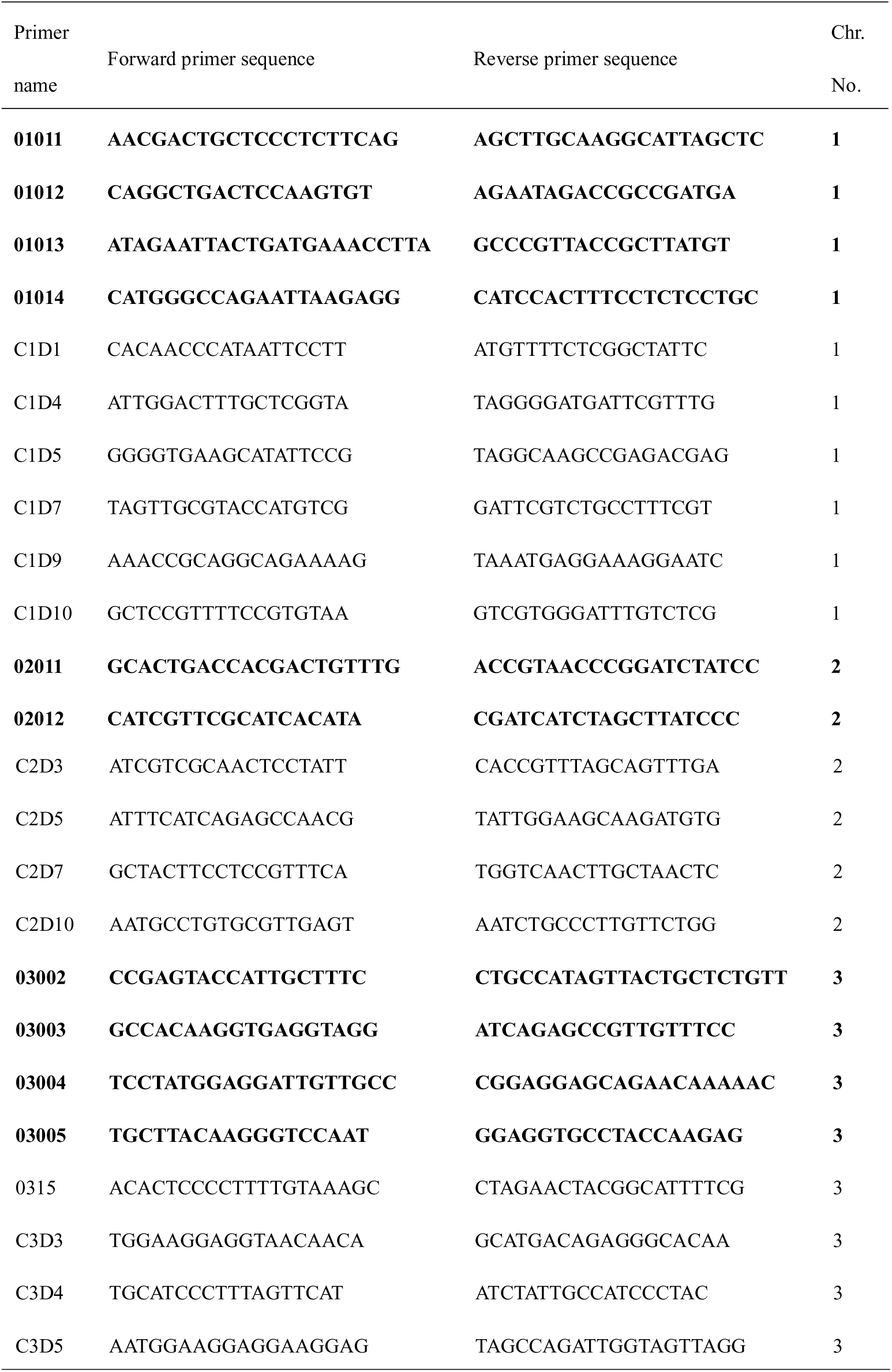
Sequences of polymorphic markers linked to *OsDMS-1* genes

To verify the true linkage relationship between the polymorphic markers and the *OsDMS-1* gene, we checked each polymorphic marker with 20 random fertile and 20 random sterile plants from the *OsDMS-1*/JH1//JH1 mapping population. The results from each polymorphic marker demonstrated that the genotype of most fertile individuals was homozygous, with the fertile parent as JH1, while the genotype of most sterile individuals was as heterozygous as the genotype of F_1_ (*OsDMS-1/JH1),* with a small number of exceptions (Fig. 3). Note that figure 3 represents the genotyping results from a single marker on each located chromosome. The fertility phenotypes track the genotypes, indicating linkage between these polymorphic markers and *OsDMS-1* sterility. These linked polymorphic markers were located on chromosomes 1, 2, and 3, which presented the possibility that three different loci control the sterility of *OsDMS-1*. This possibility is at variance with classical genetic analysis suggesting that a single gene controls the sterility of *OsDMS-1*. To characterize this potentially novel sterility mechanism, we designated the locus on chromosome 1 as *OsDMS-1A,* the locus on chromosome 2 as *OsDMS-1B,* and the locus on chromosome 3 as *OsDMS-1C*.

**Figure 3.**
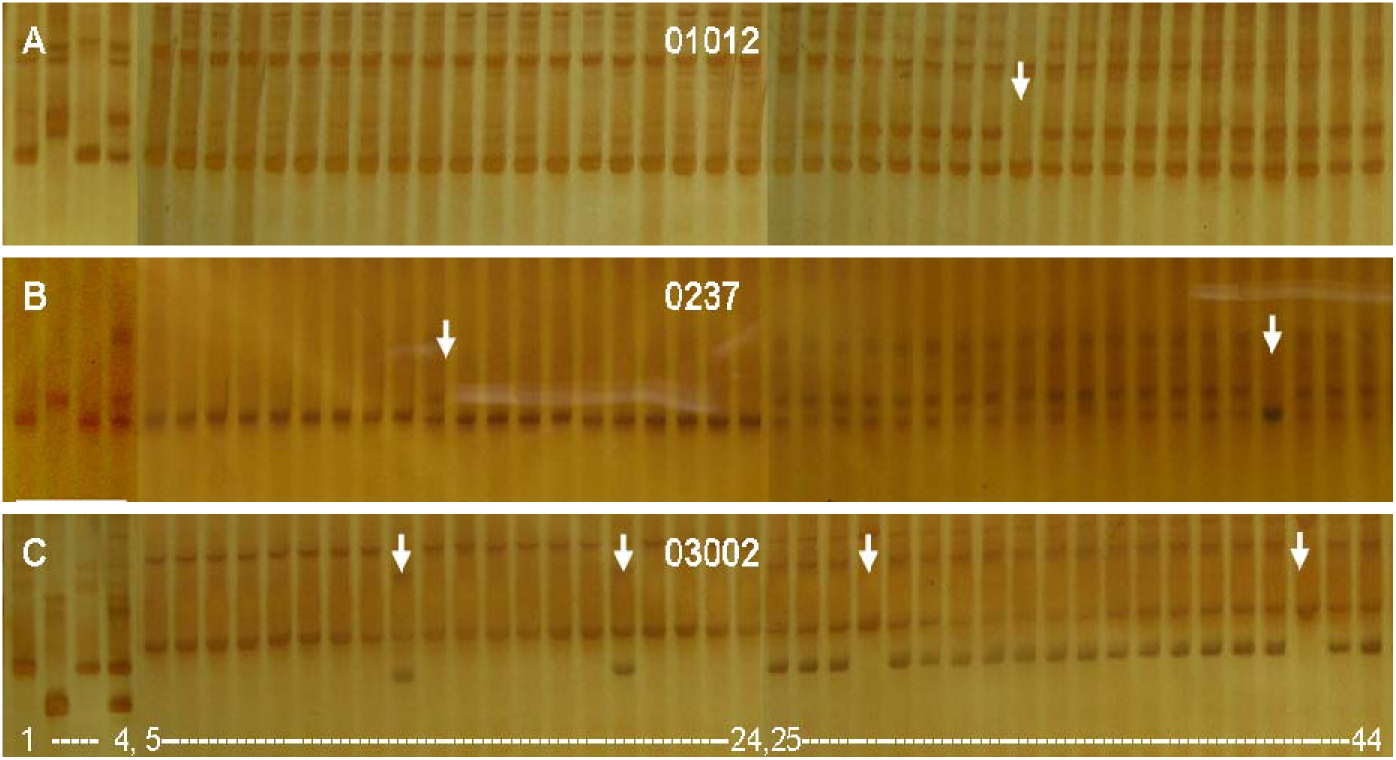
Screening and verification of linked InDel markers A-C: the genotypes of parents, bulks, and fertile and sterile individuals analyzed with InDel markers 01012, 0237, and 03002, respectively; Lane 1 represents JH1, lane 2 represents *OsDMS-1*, lane 3 represents fertile bulks, lane 4 represents sterile bulks, lanes 5-24 represent fertile individuals, and lanes 25-44 represent sterile individuals. The lanes with arrows indicate recombinant individuals.

### Determination of relative position among the loci and their linked markers, and Estimation of genetic distance

To screen for markers more closely linked to the *OsDMS-1* loci, multiple primer pairs within InDel markers on chromosome 1, 2, and 3, adjacent to our verified markers, were designated for synthesis. With these new markers, we examined an additional series of polymorphic linked markers on these chromosomes (Table 2, normal font). To map the *OsDMS-1A* locus and its linked screened markers, 344 fertile plants and 338 sterile plants from *OsDMS-1/JH1//JH1* populations were genotyped with linked markers from chromosome 1. After PCR and electrophoresis, multiple recombinants were detected by C1D1, C1D4, C1D5, C1D7, C1D9, and C1D10 on chromosome 1 (Table 3). The recombinants detected with the remaining polymorphic markers are also shown in table 3.

**Table 3.** The statistical data of the recombinants

| Primer name | Serial No. of fertile recombinants | Serial No. of Sterile recombinants | Total |
| --- | --- | --- | --- |
| C1D1 | F142 | S194 , S198 , S206 , S242 | 5 |
| C1D4 | none | S242 | 1 |
| C1D5 | none | S117 , S128 | 2 |
| C1D7 | none | S117 , S128 | 2 |
| C1D9 | F25 , F117 | S117 , S128 | 4 |
| C1D10 | F25 , F117 , F232 | S117 , S128 | 5 |
| C2D3 | F91 , F119 , 240 | none | 3 |
| C2D5 | none | none | 0 |
| C2D7 | none | none | 0 |
| C2D10 | F77 , F160 , F177, , F253 | S122 , S224 | 6 |
| 0315 | none | S72 , S176 , S179 | 3 |
| C3D3 | F211 | S154 , S191 , S201 , S223 , S237 | 6 |
| C3D4 | F109 , F147 , F187 , F211 | S154 , S191 , S198 , S201 , S223 , S237 | 10 |
| C3D5 | F109 , F147 , F187 , F211 | S122 , S154 , S191 , S198 , S201 , S223 , S237 | 11 |

Because the recombinant sets from C1D4 and C1D5/C1D7 were mutually exclusive, and the recombinants detected by C1D4 and C1D5/C1D7 were few, the locus of *OsDMS-1A* was defined between InDel markers C1D4 and C1D5/C1D7. Utilizing the recombinant numbers with the inclusion of the relationship of the recombinant sets of all markers from chromosome 1, the relative position and genetic distances of the *OsDMS-1A* locus and its linked markers were as follows: C1D1(0.73cM)-C1D4(0.15 cM)-*OsDMS-1A*-C1D5(0.30 cM)-C1D7(0.30 cM)-C1D9(0.59 cM)-C1D10(0.73 cM) (Fig. 4A). In a similar fashion, *OsDMS-1B* was localized between InDel markers C2D3 and C2D10 with genetic distances of 0.44 and 0.88cM, respectively (Fig. 4B), while *OsDMS-1C* was localized between InDel markers 0315 and C3D3 with genetic distances of 0.44 and 0.88cM, respectively (Fig. 4C). Each of these mapped loci and markers are shown in figure 4. From the results of our recombinant events survey (Table 3), we also found that the recombinant sets from chromosome 1, 2, and 3 were not mutually involved, which indicated that recombination between the locus and its linked markers on different chromosomes occurred separately. our mapping results thus indicated that there must be three loci which mutated within the same parent, Zhonghua 11, jointly controlling the sterility of *OsDMS-1*.

**Figure 4.**
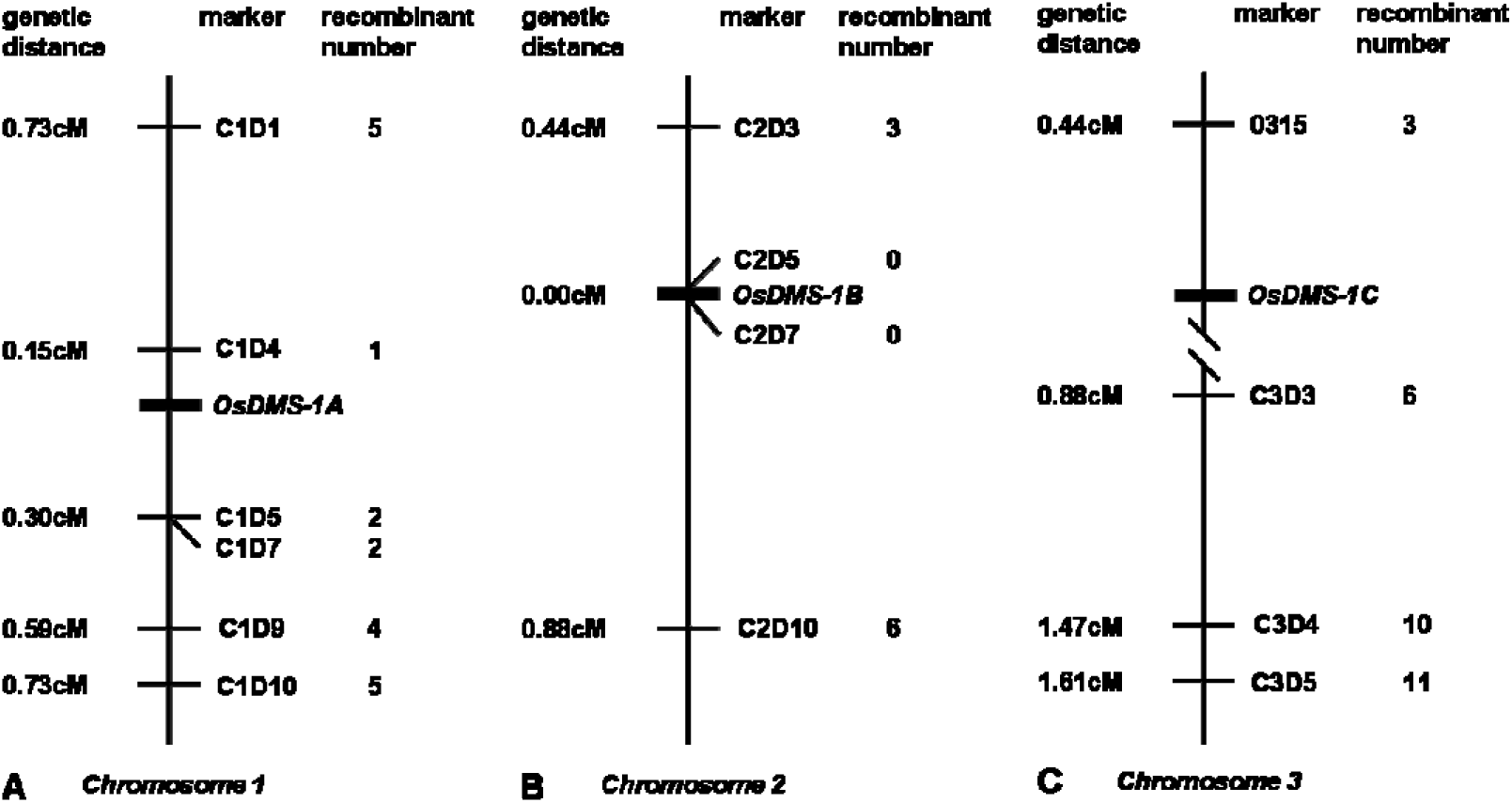
Linkage maps of *OsDMS-1* loci

## 3 Discussion

Plant dominant genetic male sterility (DGMS) represents a novel molecular male sterility mechanism. Because the restorers of most DGMS have yet to be isolated, reproduction and preservation are very difficult, with the homozygote rarely obtained. These obstacles have resulted in a paucity of DGMS data, prolonging the search for DGMS molecular mechanisms.

### *OsDMS-1* is a novel dominant male sterility mutant

Prior to our report, six rice dominant male sterility mutants had been documented: Pingxiang dominant genetic male sterile rice (*PDGMSR*) (Yan *et al*. 1989; Huang *et al*. 2007), lower temperature thermo-sensitive dominant male sterility *TMS/8987 (Deng and Zhou 1994; Li et al. 1999),* dominant genetic male sterile mutant *Zhe9248* (Shu *et al*. 2000), dominant male sterile mutants of *1783* and *1789* (Zhu and Rutger 2000), and the Sanming dominant genetic male sterility mutant *SMS* (Huang *et al*. 2008; Yang *et al*. 2012). *OsDMS-1* is a DGMS mutant found in the tissue culture offspring of a *japonica* rice, Zhonghua 11. When compared to previously reported DGMS, *OsDMS-1* displayed a different sterile phenotype and exhibited a novel mapping result. By comparison, the *TMS/8987* sterility type is a typical abortion with the sterility converting to fertility at high temperatures, and the related gene *TMS* mapped to chromosome 6 (Deng and Zhou 1994; Li *et al*. 1999); the *Zhe9248* mutant has three types of sterile pollen: typical, spherical, and stained abortion, and displays female sterility, with no gene mapping reports (Shu *et al*. 2000); the DGMS mutants of *1783* and *1789* show partial male sterility, with the ratio of selfing ~30%, with no gene mapping reports (Zhu and Rutger 2000); the Sanming dominant genetic male sterility of *SMS* is a pollenless mutant with the sterility insensitive to temperature and day length, and the *SMS* gene localized to chromosome 8 (Huang *et al*. 2008; Yang *et al*. 2012); the sterility of *Ms-p* is temperature-sensitive and can produce seed by selfing at high temperatures (Yan *et al*. 1989), with the *Ms-p* restoration gene found in six cultivars, the *Ms-p* sterility gene and the *Rfe* restoration gene were mapped to different intervals on chromosome 10 (Huang *et al*. 2007; Xue *et al*. 2009); in contrast, *OsDMS-1* displays typical sterility pollen and low numbers of stained abortion pollen, with the sterility insensitive to temperature and day length, while exhibiting female sterility. Notably, using a single BF1 population, we found that three loci on chromosomes 1, 2, and 3 were mapped and tracked the sterility of *OsDMS-1*. The novel pattern of sterility and gene mapping indicates that *OsDMS-1* is an unique dominant male sterility mutant.

### A proposed model for the sterility of *OsDMS-1*: three loci formed a triad collaboratively contributing to male sterility

Theoretically gene mapping is a reliable method to determine the genetic mechanism of dominant genetic male sterility (Huang *et al*. 2007). Most reports characterizing male sterility mutants suggest that the phenotype is controlled by one gene through classical genetic analysis, and only one gene was verified to track sterility by molecular mapping, thus, classical inheritance analysis and molecular mapping provided similar data. In our analysis of *OsDMS-1,* using an *OsDMS-1/JH1//JH1* BF_1_ population, we mapped three loci supposedly linked to *OsDMS-1* sterility. We found that all the genotypes of the fertile and sterile plants, with the exception of some recombinants, in this BF_1_ population were consistent with the phenotype of fertility and sterility. This was not an occasional genetic phenomenon as we also mapped two loci controlling dominant male sterility in another DGMS mutant of *OsDMS-2*. The mapping data not supporting the classical genetic analysis on the number of sterility loci in this report indicates a novel genetic phenomenon which can not be explained by existing models. Thus, we deduce from our data, that three dominant sterility loci must have mutated in the same Zhonghua 11 parent, with only two individual genotypes (*ABC/abc* and *abc/abc*) existing in the *OsDMS-1*/JH1//JH1 BF_1_ population. We subsequently proposed two hypotheses to explain this genetic phenomenon. We abbreviated these dominant loci *OsDMS-1A, OsDMS-1B,* and *OsDMS-1C* as *A*, *B*, and *C*, respectively; with the recessive loci as *a, b,* and *c*. respectively.

### Hypothesis one: Parent Originated Loci Tying Inheritance (POLTI)

Generally, because of the free combination of heterologous chromosomes at meiosis I, the chromosome and its loci from male and female parents are distributed into gametes randomly. Thus, a triple heterozygote such as *OsDMS-1* which has the genotype *ABC/abc,* could produce eight possible gametes, the parentals (ABC and *abc),* and each possible recombinant (*ABc, AbC, Abc, aBC, aBc,* and *abC*) with equal probability. Noting that the recurrent male parent JH1 of *OsDMS-1*/JH1//JH1 is recessive homozygous and only produce *abc* type gametes, a mechanism to explain the exclusive production of *ABC*/*abc* and *abc*/*abc* genotype individuals would postulate that at meiosis I, the fertility/sterility related loci and its chromosomes in *ABC*/*abc* F1 individuals violated the principle of random assortment, with the three different sterility loci and related chromosomes from parent *OsDMS-1* moving towards one cell of the dyad, while the three fertility loci and related chromosomes from JH1 moved towards the other cell of the dyad, resulting in the exclusive production of parental gametes; when these female gametes were fertilized by the JH1 recessive gametes, only parental genotypes would be produced in the BF1 population. This hypothesis postulates a mechanism that groups the related loci on different chromosomes from the same parent, which leads the tied heterologous loci from one parent segregating to the same gamete. We termed this model as Parent Originated Loci Tying Inheritance (POLTI).

### Hypothesis two: Loci Recombination Lethal (LRL)

The second hypothesis to explain the exclusive production of *ABC*/*abc* and *abc*/*abc* genotype individuals postulates that at meiosis I, although the three heterozygous chromosomes and the related loci comply with principle of random assortment in the *ABC/abc* F_1_ female, with all eight kinds of gametes produced at a very early stage, the recombinant gametes degenerate or abort prior to maturation, which results in no recombinant male gamete production and thus no recombinant female gamete fertilization, leading to only *ABC/abc* and *abc/abc* genotype individuals appearing in the BF_1_ population. These assumed results were in accordance with the observation of reduced pollen numbers in *OsDMS-1* anthers and a reduction in crossing seed-setting ratios of *OsDMS-1*. A variant of this model postulates that all the eight gametes develop normally and produce zygotes, or even seed, or even seedlings of all genotypes, with the recombinant offspring failing to reach maturity, causing only the fertilization products of parental gametes to survive, as was reflected in our genetic analysis and gene mapping. This possibility is remote as we failed to find obvious abortion of developing seed, low seed germination rates, or extensive seedling death. Despite these findings, we termed loci recombination resulting in death of recombined gametes or offspring produced by recombinant gametes as Loci Recombination Lethal (LRL).

These hypotheses may not reflect the mechanism producing the disagreement between classical genetic analysis and the molecular mapping of DGMS in the *OsDMS-1* mutant. In order to determine which of these mechanisms is operating in *OsDMS-1,* functional characterization of the *OsDMS-1* sterility loci is required.

## Author contribution statement

Y. Z. conceived the experiment. K. Y., B. Z., Y. C., M. S., X. C. and H. M. had the equal contributions to performing the research. Y. Z. and J. L. produced the materials used. Y. Z., K. Y. and R. C. wrote the manuscript. All authors reviewed and approved this submission.

## Compliance with ethical standards

### Ethical standards

The authors declare that this study complies with the current laws of the countries in which the experiments were performed

## Acknowledgments

This work was supported by funds from the National Natural Science Foundation of China (30970274, 31370349; http://www.nsfc.gov.cn), and the Fundamental Research Funds for the Central Universities (XDJK2015A014; http://kjc.swu.edu.cn/kxjsc/index.php).

